# Single-cell phenotypic analysis and multiplet detection through incorporation of microscopy data into cellenONE-based single-cell proteomic data analysis

**DOI:** 10.64898/2025.12.01.691577

**Authors:** Markella Loi, Stephen Holmes, ShuXian Zhou, Marie Held, Philip Brownridge, Jessica Coomber, Leandro X. Neves, Trevor R. Sweeney, Edward Emmott

**Author notes:** These authors contributed equally (Markella Loi, Stephen Holmes).

## Abstract

The reliability of single-cell proteomics (SCP) is intrinsically linked to the fidelity of cell isolation; however, the identification of co-isolated cells (doublets or multiplets) remains a persistent challenge for single-cell (sc-) omics. While single-cell transcriptomics has established probabilistic frameworks for doublet detection, these methods are ill-suited for the sparser throughput of SCP datasets. This work presents *scpImaging*, a novel computational pipeline that repurposes the latent microscopy data routinely generated during cellenONE-based sample preparation to provide deterministic quality control (QC) and high-content phenotypic profiling. We demonstrate that standard proteomic quality metrics for SCP (e.g., peptide count, signal intensity) paradoxically favour retaining doublets. In contrast, *scpImaging* applies machine-learning-based cell segmentation (Cellpose-SAM) to identify and exclude these artefacts with high precision. Furthermore, the framework integrates morphological metrics generated using the open-source CellProfiler, with proteomic abundance data, enabling joint analysis that links cell shape and texture to molecular cell state. Provided as an open-source R package, *scpImaging* offers a scalable, automated solution for enhancing data integrity and adding a phenotypic dimension to SCP experiments without increasing experimental cost or complexity.

## Introduction

Multicellular organisms function as intricate cellular systems composed of diverse cell types that collectively coordinate biochemical networks essential for growth, development, and survival^1^. Although these cells act in concert as a unified biological entity, variability in their activity and behaviour inevitably arises from stochastic molecular fluctuations and dynamic intercellular interactions^2^.

This diversity or cellular heterogeneity, is a crucial aspect of multicellular life, spanning multiple dimensions ^3, 4^. Heterogeneity is evident not only in molecular signatures, but also in phenotypic traits like morphology, cell size, or organelle architecture, each shaped by and reflecting distinct molecular pathways^5^.

Yet, conventional population-level analyses often assume cellular uniformity, thereby obscuring a fundamental aspect of biology^1^. Recognising and resolving cellular heterogeneity has thus become necessary to understand how individual cell states collectively define system-level function. To capture these unique molecular signatures, advanced analytical tools for single-cell resolution have emerged^6^. This paradigm shift began with sc-genomics and transcriptomics and has since expanded to proteomics, metabolomics, and lipidomics, collectively referred to as multiomics^7^.

SCP has witnessed remarkable progress with the advent of methods like SCoPE-MS (Single Cell Proteomics by Mass Spectrometry) and its successor, SCoPE2 (Single Cell ProtEomics) ^8, 9^. Despite this technological bloom, a shared challenge across sc-omics persists, the accurate isolation of individual, intact cells. Ideally, every isolated cell would be a singlet; however, in practice, two (doublet) or more (multiplet) cells may be inadvertently captured together in a single droplet^10^. These events can arise from cell aggregation or co-encapsulation during sample preparation, resulting in a non-negligible rate of doublets/multiplets ^11^. These artefacts can mislead clustering, inflate estimates of cellular diversity, and distort biological interpretations ^12^. Thus, rigorous QC is essential to ensure the integrity of single-cell data.

Existing QC pipelines are almost exclusively designed for sc-RNA sequencing (scRNA-seq) data. Experimental approaches such as Cell Hashing, MULTI-seq data and demuxlet introduce molecular barcodes or exploit genetic variation to distinguish singlets from multiplets^13, 14^, while computational alternatives such as DoubletFinder and Scrublet detect multiplets directly from scRNA-seq data by inferring hybrid transcriptional profiles^15^. These tools provide a cost-effective, post-hoc solution, yet remain transcriptomics-biased and do not easily translate to other omics modalities. This gap is particularly problematic in SCP, where datasets are typically smaller than those available for testing scRNA-seq approaches^16^.

A promising strategy to address this limitation involves image-based QC, utilising imaging data to evaluate the physical and morphological integrity of individual cells prior to downstream molecular analysis. This would allow a direct, modality-independent assessment of cell quality, enabling the identification of doublets or other artefacts that may compromise data interpretation. By integrating visual information into proteomic workflows, this approach would also introduce a new dimension of analytical robustness and biological insight. Visually Aided Morpho-Phenotyping Image Recognition (VAMPIRE), a sc-transcriptomics pipeline, illustrates the feasibility and analytical value of incorporating image-derived metadata in single-cell analysis – reinforcing the idea that they can be effectively used to assess cell quality^5^. However, implementations like this remain tightly coupled to specific instrument ecosystems, leaving no open or broadly deployable counterparts suited for single-cell proteomic workflows.

Automated systems such as the cellenONE with its embedded imaging offers a natural point and a technically compatible environment in which image-based QC could be adapted and extended to the requirements of SCP. Designed for high-precision single-cell sample preparation, the cellenONE combines acoustic droplet ejection with nanolitre-scale liquid handling to achieve accurate and reproducible single-cell deposition^17^. Crucially, the system operates in both transmission and fluorescence imaging modes, capturing images of every isolated cell in real time^18^. These images, routinely generated but typically discarded, constitute a rich, untapped metadata source that can be repurposed for morphological QC.

Building on this principle, we developed *scpImaging*, a computational pipeline that exploits cellenONE-derived images for image-based QC in SCP. The module integrates three main computational environments, beginning with Cellpose for deep-learning–based segmentation and annotation of individual cells^19^; CellProfiler to extract quantitative morphological and textural features^20^; and RStudio for preprocessing, data handling, and visualisation using a custom R package: ‘scpImaging’ and the ‘iSEE’ packages^21^.

By incorporating these image-derived phenotypic metrics into the SCP workflow, *scpImaging* establishes a framework that unites morphological and molecular profiling at single-cell resolution. This integration enables a more comprehensive characterisation of cellular heterogeneity, offering not only improved data quality but also deeper insights into how cell morphology and proteomic signatures jointly might define cell state and function.

This workflow represents, to our knowledge, the first SCP pipeline capable of automated doublet detection using existing metadata collected by the cellenONE during sample preparation for QC, without adding experimental burden and simultaneously integrating a phenotypic layer into SCP datasets – building a promising basis for single-cell multi-modal analysis. Importantly, these QC and phenotypic assessments are independent of downstream applications and are equally applicable to any cellenONE-based omics, or even cell line development uses employing the cellenONE instrument.

## Materials & Methods

### Cell Culture

HEK-293T (Hiscox Lab, Liverpool), A549-ACE2-TMPRSS2 (Pasteur Institute, France), Huh7 (Davidson Lab, Bristol) and BV-2 (Goodfellow Lab, Cambridge) cells were cultured in Dulbecco’s Modified Eagle’s Medium (DMEM) supplemented with 10% foetal bovine serum (FBS), 1% non-essential amino acids (NEAA) and 1% penicillin/streptomycin (P/S). Cell lines HTB-40 (ATCC) and CRFK (ATCC) were grown and preserved in Minimum Essential Media (MEM) completed with 1% L-glutamine in addition to the aforementioned supplements. All cells were maintained at 37°C with 5% CO_2_ and tested monthly for mycoplasma.

### Generation of carrier and reference

Cells were mixed in a 50:50% ratio (HEK-293T: A549-ACE2-TMPRSS2) to generate bulk lysate. Cells were resuspended in LC-MS grade water to a final concentration of 4000 cells/µl and were then probe sonicated for 6 x 10 seconds pulses at 30% amplitude, with 30 second rest intervals on ice. A total of 320,000 cells worth of material was combined with anhydrous dimethyl sulfoxide (DMSO, 40% final), 500mM HEPES (2-[4-(2-hydroxyethyl)piperazin-1-yl]ethanesulfonic acid) containing 0.04% n-dodecyl-β-D-maltoside (DDM), and 2µg of gold standard trypsin (11.1ng/µl final). Following a brief centrifugation to collect all the liquid, samples were digested at 37°C, 1,300 rpm for 16 hours. Two aliquots of approximately 70,000 cells were then labelled using 0.333µg/µl of either the 126C or 127N labels of an 18plex tandem mass tag (TMTpro) batch (Initial HEK-293T/A549 dataset 126-134N, LOT: ZD390445 and134C-135N, LOT: ZA381335. Singlet/Doublet HEK-293T/A549 dataset 126-134N, LOT: VK311069, 134C-135N LOT: XC343801). Cells were incubated in a thermomixer for 1 hour at 21°C, 1300 rpm to allow labelling. Samples were quenched using hydroxylamine (HA, 0.045% final) and further incubated in for 30 mins at 21°C, 1300 rpm. The carrier (126C) and reference (127N) labelled samples were then combined in a 40:1 ratio to obtain a final cell ratio of 200:5 (Carrier: Reference). These samples were then dried down and resuspended in 0.01% DDM in 0.1% formic acid (FA) to obtain a stock of carrier and reference at 100 cell carrier: 2.5 cell reference per µl. Of this, 2µl was added to each required well for cellenONE pickup.

### CellenONE

Samples for SCP were prepared according to the nPOP v2 method^22^. Type 2 piezo-dispensing capillaries (PDCs) were used for cell isolation, reagent dispensing, and pickup. LC-MS grade DMSO was first dispensed onto SciCHIP H1 coated glass slides in predefined multiplexing layouts (14-plex). Follow-up scans of the dispensed droplets were used for quality checks, and the images were examined for any off-target droplets. Cells of interest were harvested using Trypsin-EDTA (1X). The derived pellets were washed with phosphate-buffered saline (PBS) and resuspended in PBS to a final concentration of 200 cells/μl. Cell suspensions prepared for isolation were initially mapped using the instrument’s cell-sorting mode. Single cells were finally isolated based on cell size and shape – these parameters were specific to each cell line. Each isolated single cell was dispensed onto DMSO-patterned regions of the SciCHIP slides.

A digestion master mix was prepared immediately before use consisting of 10mM HEPES and 0.05% DDM in LC-MS grade water. The solution was placed within an insulated ice box and degassed for 20 minutes. Trypsin gold (100ng/µl) was added, and the mix was further degassed for an additional 10-minutes.

The mastermix was then dispensed onto each cell-containing droplet, and the slides were incubated overnight at room temperature to allow complete proteolysis. TMTpro labelling was performed the following day with labels 128C-135N. Single-cell samples were retrieved into 384-well plates. In the collection wells, 0.01% DDM was included to reduce peptide adhesion to plastic surfaces^23^. Samples were collected from the glass slide to the 384-well plate in 50:50 acetonitrile/ LC-MS grade water. Following collection, samples were dried in a speedvac, sealed and stored at −80 °C until required.

### LC-MS/MS

Analysis by LC-MS/MS was carried out on a Dionex Ultimate 3000 coupled to a Q-Exactive-HF mass spectrometer. The system was configured for direct injection to the analytical column (Aurora® 3 Ultimate™ 25×75 XT C18 UHPLC column). The LC-system uses buffer A (0.1% formic acid in LC-MS grade water) and buffer B (0.1% formic acid in LC-MS grade acetonitrile, 99.9%). The column is first equilibrated using 3.2% buffer B for 11 mins. Peptides are loaded in 3% acetonitrile and 0.1% trifluoroacetic acid (TFA) and then eluted in a linear gradient of 6.4-20% of buffer B over 45 mins at 200 nl per min. Immediately followed by a washing step at 95% buffer B for 3 mins. The column is then equilibrated at 62 mins for a duration of 13 mins at 3.2% buffer B. The Q-Exactive-HF was operated in data dependent acquisition mode, with survey scan acquired at a resolution of 120,000 at 200 m/z over a scan range of 450-1600 m/z. Top 7 most abundant with charge states +2, +3 or +4 were selected for fragmentation and MS2 analysis at 120,000 resolution with an isolation window of 0.7 m/z with an NCE of 31%. The maximum injection time for both MS1 was 100ms with a maximum AGC target of 1e6, while MS2 was 300 ms, with a maximum AGC target of 5e4. Dynamic exclusion (30.0s) was enabled.

### LC-MS/MS Data analysis

Raw mass spectrometry data were processed using MaxQuant (version 2.7.5) with the reporter MS2 search mode enabled for 18-plex TMT quantification ^24^. TMT correction factors corresponding to the lots used (see above) were applied during quantification.

Carbamidomethylation was not included as a fixed modification. Variable modifications were set to oxidation (M) and acetylation (protein N-terminus). The acquired spectra were searched against the UniProt human reference proteome (UP000005640 – accessed 28^th^ August 2025, 20,659 entries), with trypsin set as the proteolytic enzyme and missed cleavages set at 2. The option “Calculate peak properties” was enabled. False discovery rate (FDR) thresholds were set to 1% at both peptide and protein levels. Data-driven Alignment of Retention Times for IDentification tool (DART-ID) was used to update PEP values by incorporating retention time information as previously described^25^.

### Cell segmentation (Cellpose)

Cellpose-SAM was used for cell segmentation^26^. Pre-processed micrographs, representing 75 x 75 pixel cropped cellenONE micrographs were subject to manual or automated segmentation in Cellpose-SAM (version v3.1+). To establish an automated model, multiple cell lines (A549/ HEK-293T/ HTB-40/ Huh7/ BV-2/ CRFK) were isolated with the cellenONE as described above. To ensure technical variation, a PDC was used for each biological replicate, in addition to preparing each cell line in triplicate – specifically 220 cells per cell line, per replicate. The acquired micrographs were randomised and split into 70% training (2772 cells) and 30% (1188 cells) validation subsets. Three independent annotators manually segmented the sets within Cellpose-SAM to create high-fidelity reference masks. The training data were used to train a custom Cellpose-SAM model for automated cell segmentation from cellenONE micrographs, available through Figshare.

### Cell profiling (CellProfiler)

An analysis pipeline was established for Cellprofiler^20^ (version 4.2.8), which also produces a project file, suitable for loading into Cellprofiler analyst^27^ for further image analysis and classification within Cellprofiler if desired. The pipeline employs the “measureimagequality”, “measuregranularity”, “measureimageareaoccupied”, “measureimageintensity”, “measureobjectintensity", “measureobjectintensitydistribution”, “measureobjectsizeshape”, and “measuretexture” modules, and exports to spreadsheet, database, and saves images. A copy of the pipeline file is available on Figshare.

### Cell status and phenotype data analysis

#### Data analysis was conducted in R (4.4.3)

Single-cell proteomic analysis was conducted using the bioconductor ‘scp’ package, and its linear-modelling-based *scplainer* workflow^16, 28, 29, 30^. Sample annotation tables for data annotation were generated using our in-house software *scpAnnotator* (Holmes et al. In preparation). In brief, zeros were converted to NA, the sample to carrier ratio was calculated, and samples were filtered based on preimplantation factor (PIF), DART-ID “qvalue” and the sample to carrier ratio. Next the number of observed peptides, median intensity, and median coefficient of variation (CV) per cell were calculated and cells filtered. Peptide spectrum matches (PSMs) were then aggregated to the peptide level. Peptide results from individual experiments were aggregated to a single experiment, and the peptide data log_2_ transform log_2_ peptide data was then used for linear modelling using the *scplainer* approach, with the model accounting for median intensity, TMTpro channel, LC-MS/MS run, and either “Cell type” or “Doublet status”, depending on the analysis. ANOVA-principal component analysis (APCA) was used for principal component analysis of the data.

Analysis of Cellpose-SAM accuracy relative to human annotators was performed using built-in Cellpose functions, via the reticulate R package. Custom R functions were used for image cropping, overlaying masks, counting cells, and appending image metadata to the “singlecellexperiment” object or “dataframe.”

Morphology-related metrics extracted via CellProfiler, were evaluated for their discriminatory ability using three classifiers: a general linear model (GLM), a random forest model and a support vector machine approach (SVM). Classifier performance was assessed by receiver operating characteristic (ROC) curves, plotting true positives versus false positive rates across varying thresholds.

Finally, interactive data visualisation was performed using the ‘iSEE’ package^21^ and a custom panel for cellenONE micrograph visualisation. Custom functions for image processing and linking image metadata to “singlecellexperiment”/ “QFeatures” or “data.frame” objects, and the custom cellenONE iSEE panel are available through the ‘scpImaging’ R package.

### Data and model availability

The proteomics data, and associated metadata are available through the EBI PRIDE repository, accession numbers PXD071380 (HEK-293T/A549 dataset), PXD071269 (HEK-293T/A549 Singlet/Doublet dataset). CellenONE images for model training and validation on multiple cell lines (10.6084/m9.figshare.30724478), the Cellpose-SAM model (10.6084/m9.figshare.30685490) and CellProfiler pipeline (10.6084/m9.figshare.30685550) are available through Figshare.

The R scripts used for data analysis and production of the figures in this manuscript are available from Github at https://github.com/emmottlab/scpImagingPaper_2025. The ‘scpImaging’ R package containing the necessary functions to apply this workflow to your own cellenONE-based single-cell or SCP workflows and visualise cellenONE images using the ‘iSEE’ bioconductor app, is available from Github https://github.com/emmottlab/scpImaging.

The SCP dataset from this manuscript can be interactively explored an iSEE-based^21^ shiny app available at https://emmottlab.shinyapps.io/scpImaging_HEKA549/, and the code for this shiny app is available from Github at https://github.com/emmottlab/scpImaging_HEKA549_iSEE/.

## Results

### Failed cell dispensing, doublets and multiplets can be identified at low frequency in cellenONE-based single-cell proteomics data

To investigate doublet capture rate in cellenONE-based SCP, two distinct cell populations, A549 and HEK-293T, were resolved. The latter are human embryonic kidney cells and represent one of the most used cell lines across omics research due to their extensive characterisation^31^. A549 cells are a well-characterised human lung adenocarcinoma line, frequently used in influenza studies^32^. These two lines serve as biologically distinct single-cell systems for benchmarking our workflow.

Here, we show with an exemplar HEK-293T/A549 two cell line SCP dataset with evident doublets as identified in cellenONE-based sample preparations from the micrographs recorded by the cellenONE. The data analysis was conducted using a linear modelling approach, without imputation using the ‘scp’ bioconductor package^28^. Dataset filtering steps are shown in **Fig. S1**, and dataset QC in **Fig. S2**. Missing value reporting, as per community guidelines^16^ is given in **Fig. S3**. In the initial dataset, 6 doublets were identified (**Fig. 1**), representing less than 1% of the sorted cells in the dataset. Many of these doublets are subsequently filtered out using standard metrics like CV and median intensity. However, at least one doublet makes it into the final dataset. Although rare, such artefacts can evidently persist in SCP workflows and should be carefully monitored.

**Figure 1:**
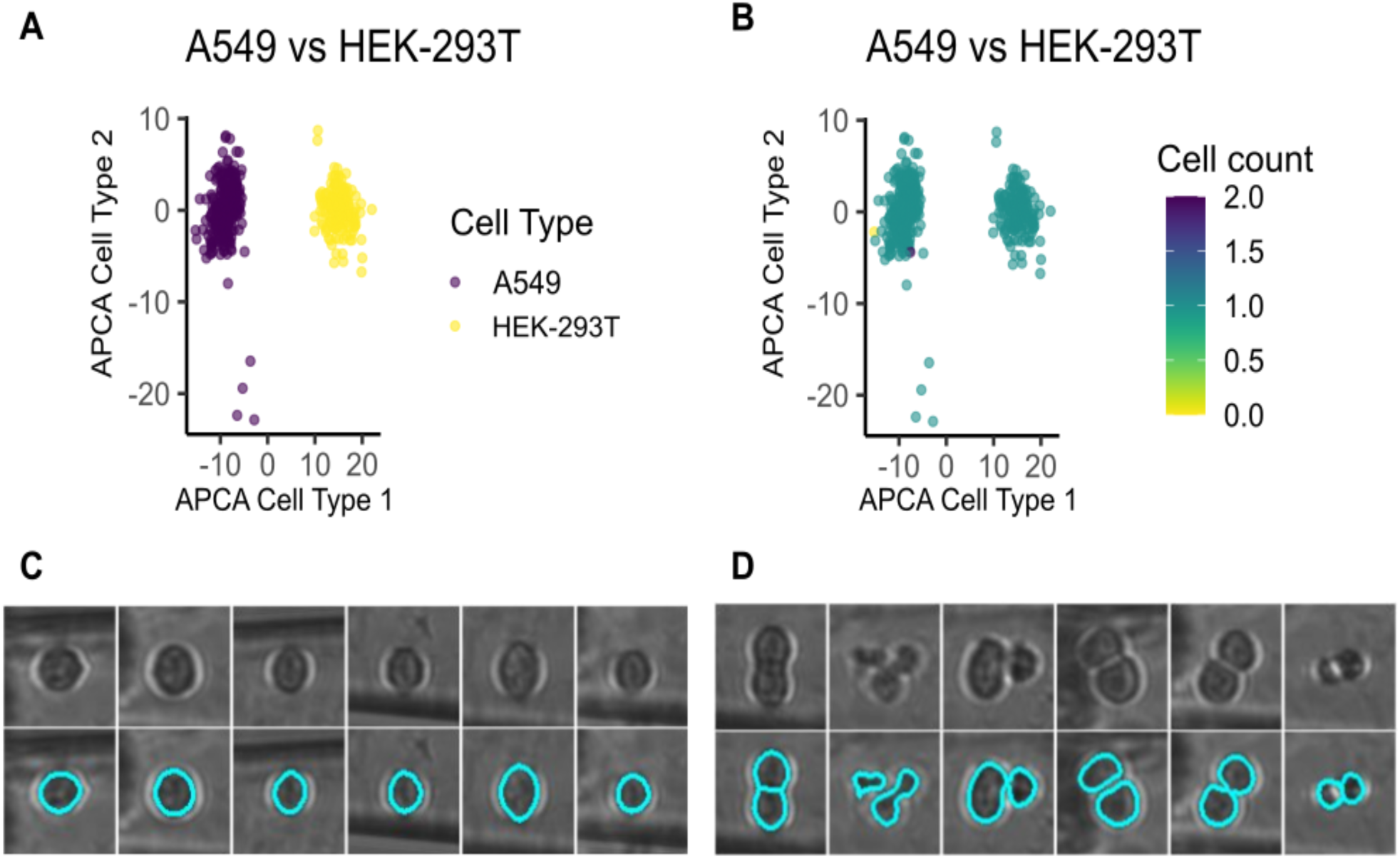
Doublets in single-cell proteomic samples. PCA of a HEK-293T/A549 SCP dataset, coloured by A) cell type, and B) cell count. Manual inspection of images identifies both C) singlets, and D) a small population of isolated doublets. Image pairs show the cropped micrographs (upper) and with the segmented cells outlined in blue (lower).

### A custom Cellpose-SAM model efficiently identifies cells in cellenONE images at scale, at an accuracy comparable to or exceeding manual segmentation

Manual image segmentation is possible, and in our hands, typical manual segmentation permits annotation of approximately 400 cropped single-cell images per hour. While this is sufficient for modest-scale SCP studies, automated segmentation offers a potential route for reliable segmentation of larger numbers of cells. We evaluated the newly released Cellpose-SA (Segment Anything Model) for image segmentation of cellenONE micrographs^26^. A Cellpose-SAM model was custom-trained on a curated dataset spanning six mammalian cell lines of different origins and sizes (HEK-293T, HTB-40, A549, Huh-7, CRFK, and BV-2). Samples were prepared in biological triplicates, isolating 200 cells per cell line replicate, using a different PDC each time. This dataset was manually annotated by three independent annotators, and the images were randomly assigned into a training dataset (70% of images) and a validation dataset (30% of images). The customised model was trained on the training dataset for 100 epochs, using the annotator one training image set **(Fig. 2)**.

**Figure 2:**
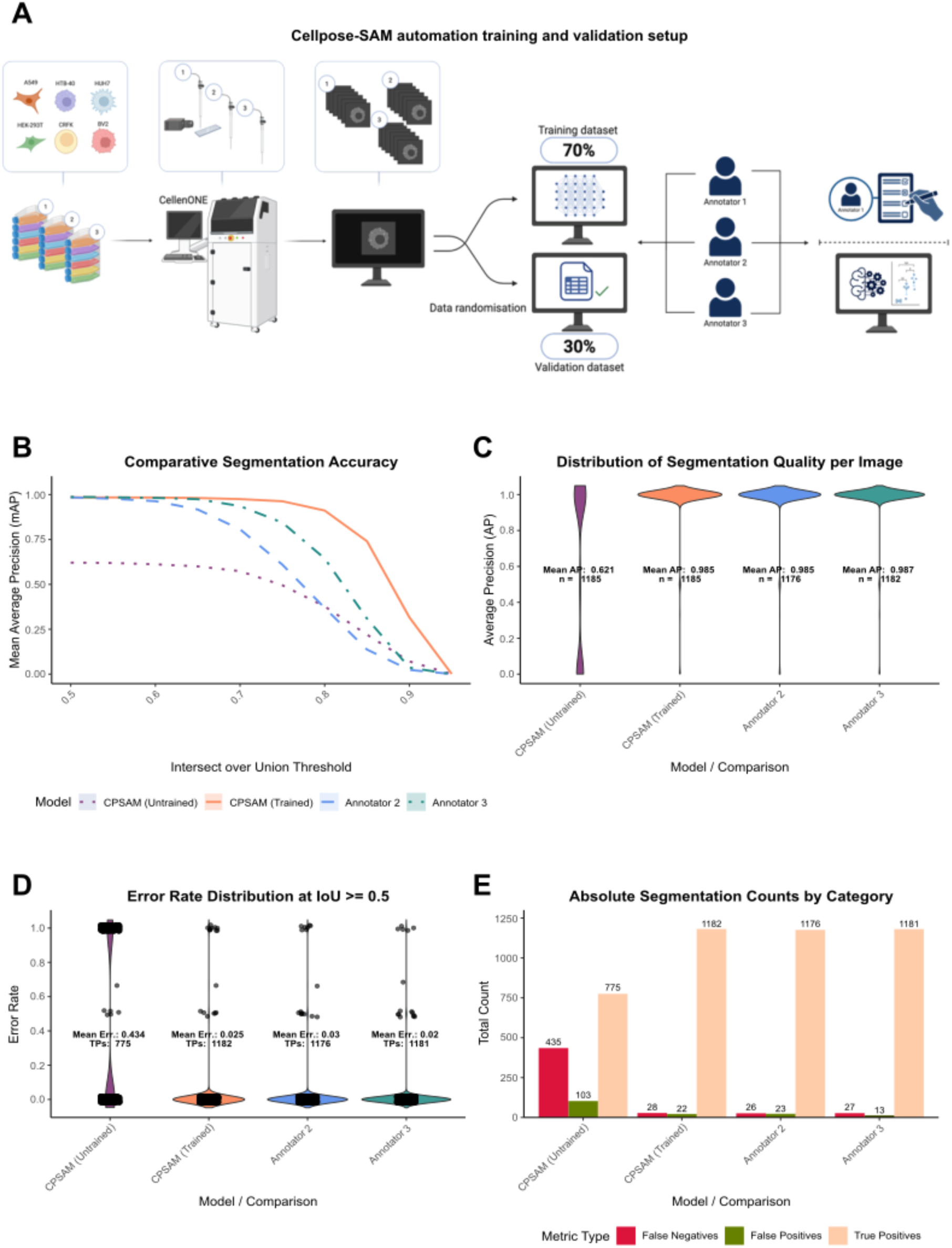
Cellpose model performance across cell lines and different piezoelectric capillaries used for cell dispensing. Panels compare default untrained and trained Cellpose-SAM models, and human annotators with human annotator 1. A) Cellpose training and validation dataset: six cell lines were used, in three biological replicates and dispensed on the cellenONE instrument. Micrographs obtained were split into training (70%) and validation (30%) datasets and manually annotated by three independent annotators. The training data from annotator 1 were used to train a custom Cellpose-SAM model. For parts B-E, all comparisons are relative to annotator 1. B) Segmentation accuracy comparing the average precision at different intersection over union (IoU) thresholds for image masks. C) Distribution of Segmentation quality scores. D) Distribution of error rates. E) Absolute segmentation counts split between false negatives, false positives and true positives.

**Figure 3:**
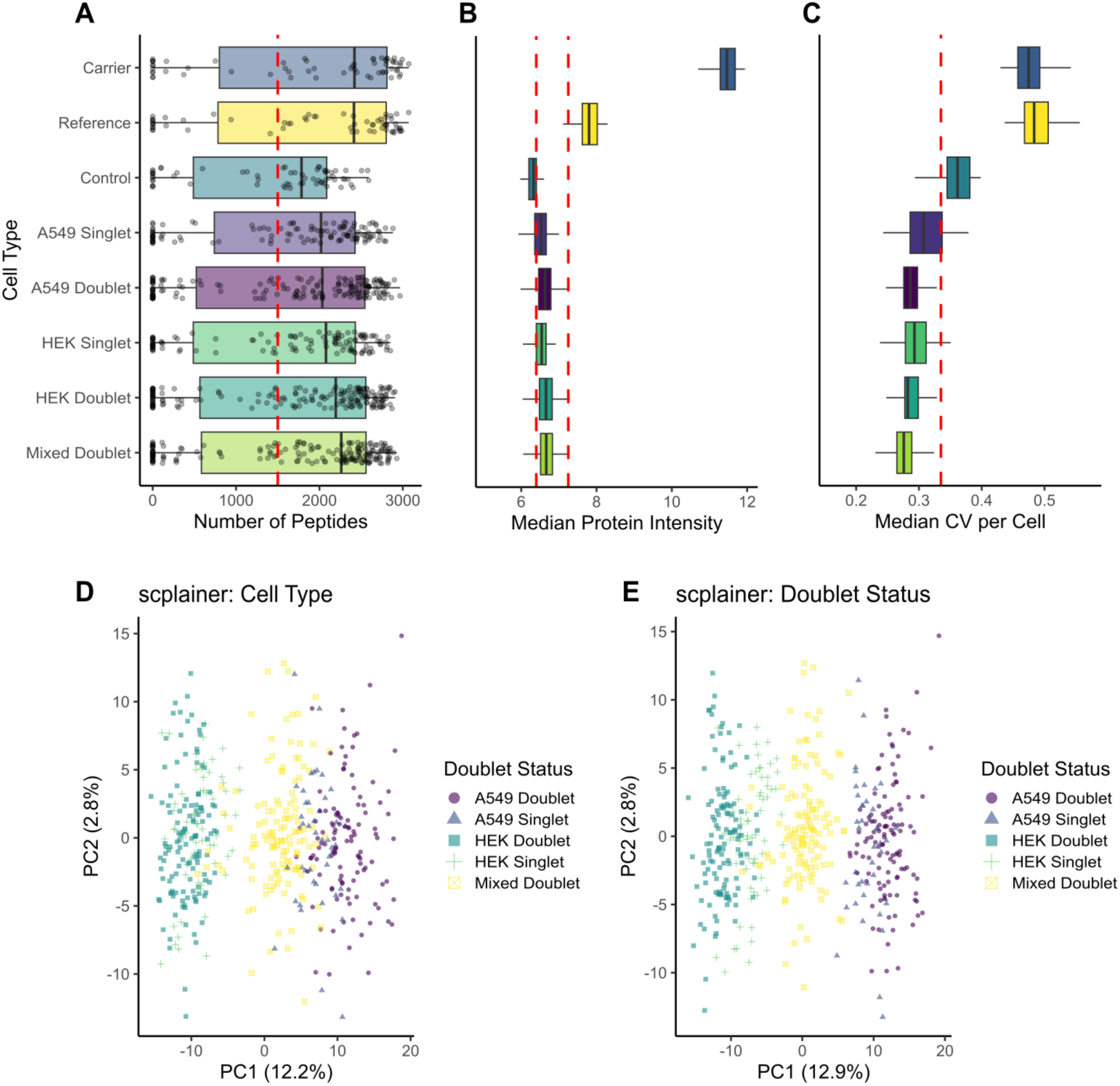
Examining doublet features in a ground-truth dataset identifies features of homo- and heterotypic doublets in single-cell proteomics data, and confirms that standard SCP-filtering fails to remove doublets. Panels A-C show metrics typically used for identifying and filtering low-quality single-cells from datasets: A) The number of identified peptides, B) Median protein intensity, and C) Median Coefficient of Variant per Cell. D and E show principal component analysis of A549 and HEK-293T dataset with doublets. Scplainer-based analysis was used, with the linear modelling incorporating either D) Cell type (A549 and HEK-293T) or E) The Singlet/doublet status of individual cells.

On the default configuration Cellpose-SAM model, the software performs poorly on cellenONE micrographs **(Fig. 2**) compared to the manual annotators. When comparing false negatives this is especially apparent with the untrained model showing poor overlap with the annotator masks **(Fig. 2B)** and failing to detect over a third of the annotated cells (**Fig. 2E**). In contrast, the trained Cellpose-SAM model performed on a par with the human annotators in terms of false positive and false-negative identifications **(Fig. 2E)** and exceeded annotators 2 and 3 in terms of cell mask overlap **(Fig. 2B).** These patterns were also observed when comparing the average precision **(Fig 2C**) and error rates **(Fig. 2D**) of both models with the human annotators. Preliminary testing also investigated model performance with the models available with earlier versions of the Cellpose software (e.g. Cyto3), these gave inferior results to the Cellpose-SAM model and were not investigated further (data not shown). Ultimately, training the module has substantially improved its performance across all metrics, exceeding its default version and even human annotators in certain cases, in line with the observations in the Cellpose-SAM manuscript^26^.

### Examining properties of doublets in single-cell proteomics data

As doublets appear rare in our cellenONE-based SCP data, our trained Cellpose-SAM model served as the basis to generate a ‘ground-truth’ SCP two cell line (HEK-293T/A549) dataset consisting of singlets, homotypic doublets, and heterotypic doublets -- with cell counts confirmed by microscopy. To construct this dataset, we deliberately designed a multiplex layout as part of the nPOP sample preparation protocol wherein individual DMSO droplets contain 0 (control), 1 (singlet), or 2 (doublet) cells. This allowed us to generate known homotypic (e.g. HEK-293T and HEK-293T) or heterotypic (HEK-293T and A549) doublets. Dataset QC and reporting metrics are shown in **Fig. S4-S6**.

Upon screening, it is clear that the standard filtering steps performed as part of minimal single-cell proteomic data processing (e.g. filtering on number of peptides, CV, median intensity) fail to remove doublets. Indeed, doublets score slightly better on these metrics compared to the singlets, as these screening steps are designed to exclude cells with low-quality quantitative data. This is a rather expected finding, as we would predict the higher protein level expected to be present in doublets relative to singlets would result in higher signal, and therefore stronger quantitative and qualitative performance based on these metrics.

The data separate clearly along the first principal component with equivalent results whether the *scplainer* model^28^ defined accounts for cell type, or the doublet status/cell identity specified. Both HEK-293T and A549 cells separate along PC1, with the heterotypic doublets forming an intermediate population, as expected, and has been observed in scRNAseq data^15^. The population is located slightly closer to the A549 singlet population, which may reflect the higher cell size/average protein amount present in A549 cells relative to HEK-293T cells. Homotypic doublets, where both cells derive from the same population are more challenging to identify, since they overlap with their corresponding singlet populations. In our data, the homotypic doublets shift to the periphery of the A549 and HEK-293T singlet populations, occupying the extremes of the PC1 distribution. This phenotype is retained when the *scplainer* model specifies either cell type or doublet status, though is clearer in the latter case.

### CellenONE micrographs can be processed to yield additional phenotypic metadata that correlates with known cell phenotypes and cell classification based on micrographs alone

In addition to recording micrographs of isolated cells, the cellenONE provides image-based cell metadata with its output files, comprising cell diameter, circularity, elongation, intensity, and where applicable fluorescence intensity.

We next sought to investigate if we could obtain further phenotypic information from the segmented cell images to complement the downstream SCP data, and expand our analyses to incorporate phenotypic data on cell morphology. To achieve this, the open-source CellProfiler software was employed to extract richer phenotypic profiles from the segmented transmitted light images, generating over 200 metrics per cell^20^. These include size metrics, capturing cell area and dimensions; shape, describing geometric properties like circularity; and texture, quantifying pixel intensity variations to estimate granularity^20^.

The cellenONE produces transmitted-light images, with the focus maintained on the capillary used for dispensing rather than on the single cells themselves, potentially leading to out-of-focus images and lower resolution. As such, we first assess if the image data was sufficient to permit cell-line classification. To evaluate the biological relevance of these measurements, the data were first normalised by robust median absolute deviation and then graded with three separate classifiers: a GLM (**Fig. 4A**), a random forest model (**Fig. 4B**) and a SVM (**Fig. 4C**). Based on the cell mask areas, all three classifiers demonstrated the ability to distinguish the cell lines on image-related metrics extract with Cellprofiler alone. Values for the area under the curve (AUC) range between 0.845-0.878. The distribution of these features across the two cell types is shown in **Fig. S7**. When permitted to use metadata from the whole image, in addition to the cell-derived metadata, the AUC values exceeded 0.95, however this amplification could be attributed to batch effects, such as sample build-up on the capillaries between these two cell lines. Consequently, we abstained from pursuing use of the whole images, as opposed to segmented cells further with this specific dataset.

**Figure 4:**
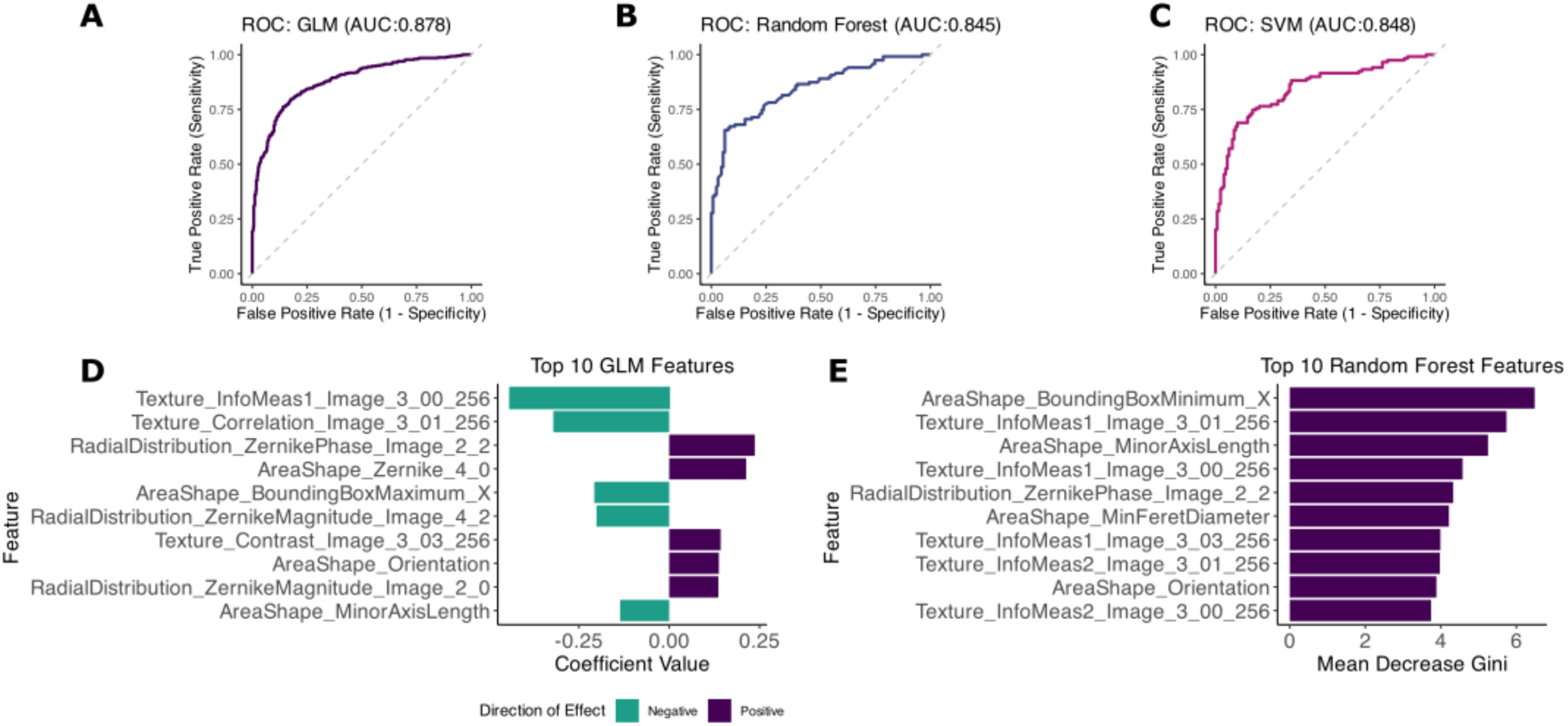
Extracted phenotypic data from cellenONE images is sufficient to permit cell-type classification. Cell phenotype data generated through Cellprofiler analysis of cellenONE HEK-293T/A549 images was robust MAD normalised and used for cell type classification. ROC curves for A) GLM, B) random forest, and C) SVM classifiers. The top image features driving the D) GLM and E) Random Forest classifiers are shown.

An advantage of both the GLM and Random Forest models is the direct output to assess which features have contributed to cell classification. Cell shape and texture appear as the primary drivers distinguishing these cell types in both models (**Fig.** 4**D****, 4E**).

### Morphological data from micrographs can be integrated into SCP analysis for a multi-modal model of single cell biology

In reviewing the correlation between our morphology data and our single-cell proteomic analysis, we assessed the significance of these associations (Pearson correlation p-values; **Fig. 5A)**. These can be overlaid onto the PCA for further data analysis as desired. We see clear, significant correlations with PC1 and PC2, and weaker correlations with other PCs **(Fig. 5A**). When overlaid onto the PCA **(Fig. 5C**), a positive correlation of “RadialDistribution_ZernikePhase_Image_8_6” with the HEK-293T cells, and negative correlation with the A549 cells is apparent (**Fig. 5D)** and vice versa when “Texture InfoMeas1_Image_3_00_256” is overlaid on the PCA. Cell texture and shape appear to have another key role in this study, as they also demonstrate the strongest positive correlation with our proteomics signatures, further highlighting their importance as significant features for multi-modal analysis.

**Figure 5:**
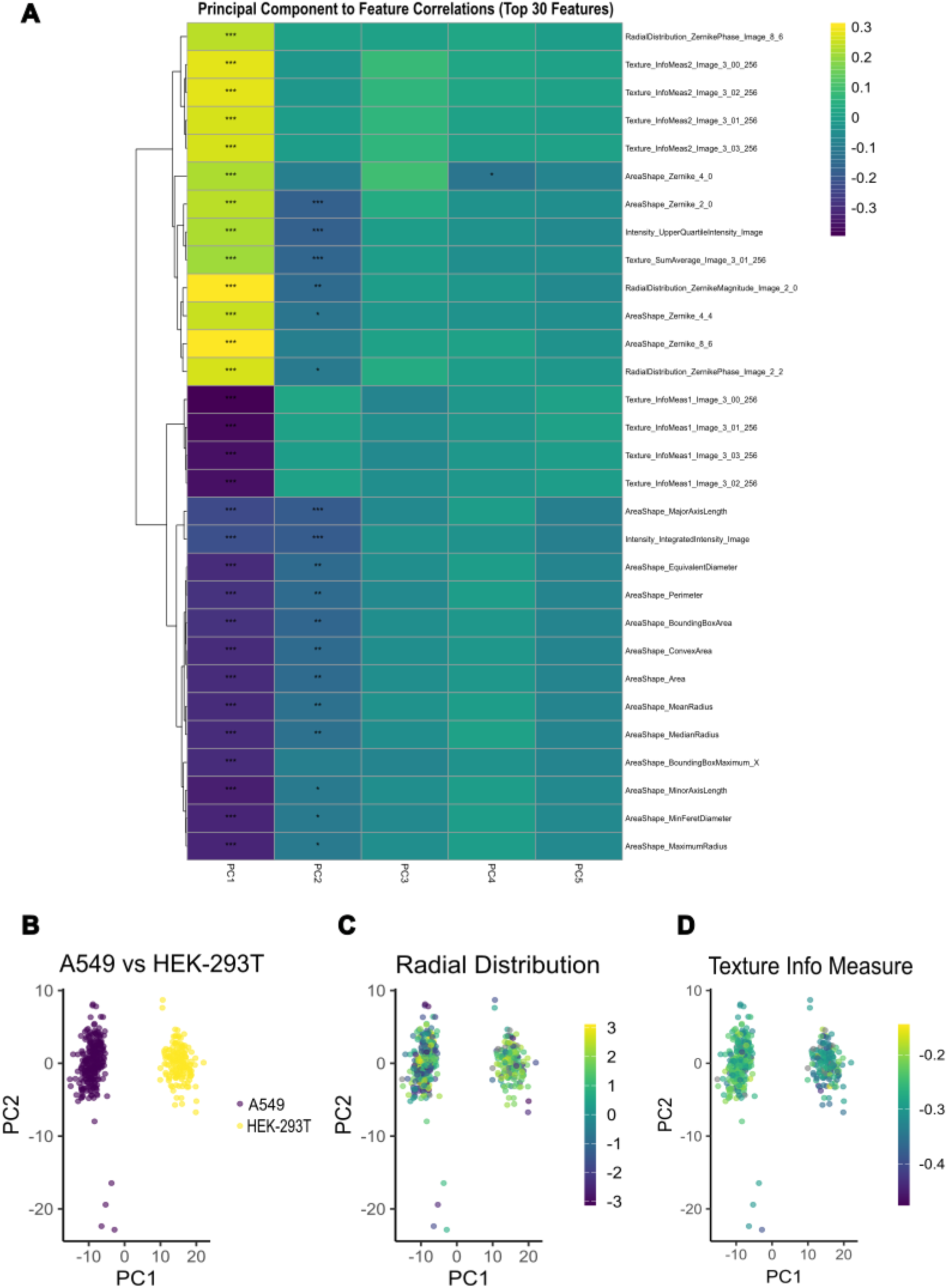
Cell morphology data correlates with single-cell proteomic data, permitting multi-omic analyses of a single-cell proteomics dataset. A) Heatmap showing the correlation of the top 30 image features with the first 5 PCs from the HEK-293T/A549 dataset from Figure 1. *, ** and *** represent Pearson correlation p-values of 0.05, 0.01 and 0.001 respectively. B-D) Principal component analyses of the same dataset coloured by B) Cell type, and the top C) positively- and D) negatively correlated image features for this dataset.

### A pipeline for processing cellenONE-based imaging data for single-cell analysis

Thus far, our findings highlighted that cellenONE micrographs can support doublet detection and also provide additional information on single cell phenotypes. Nonetheless, to assist the wider inclusion and adoption of cellenONE-based microscopy and metadata into single-cell proteomic analysis, we developed a data processing pipeline that is adaptable for different user needs **(Fig. 6).**

**Figure 6.**
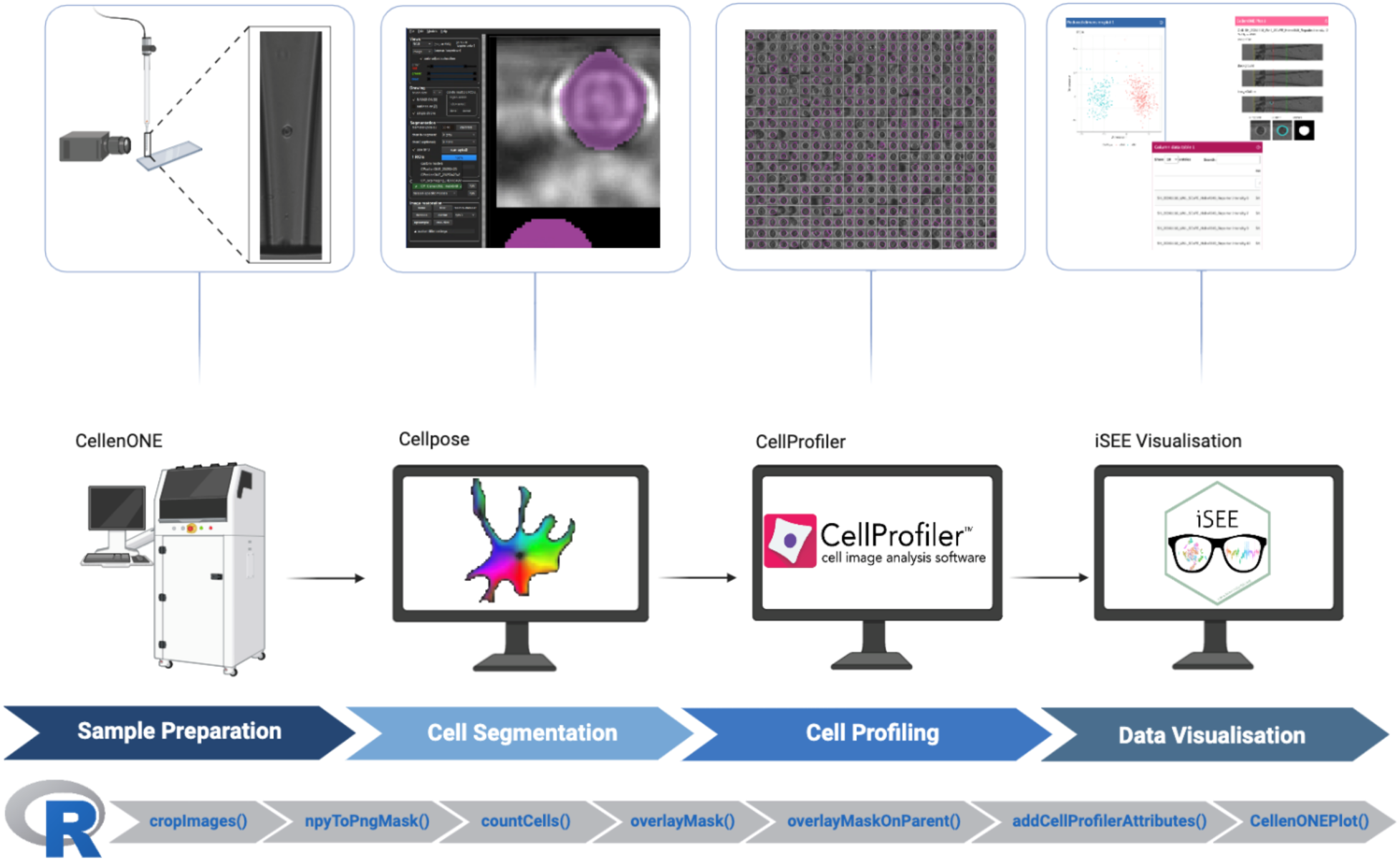
The *scpImaging* workflow. The *scpImaging* workflow incorporates several open-source tools: Cellpose-SAM for cell segmentation, CellProfiler, for extraction of morphological data from micrographs, and ‘iSEE’, for interactive data visualisation. ScpImaging is both a workflow connecting these tools, and the name of the R package that provides the necessary functions to enable this workflow.

The pipeline is modular, so it can be used for either multiplet detection, or incorporation of additional image-based metadata into downstream analysis via Cellprofiler or executing both tasks. These actions are supported by an R package: ‘scpImaging’, providing the necessary functions to enable and link the pipeline steps. This package is available from https://github.com/emmottlab/scpImaging Github.

While designed to support single-cell proteomic analysis, the *scpImaging* pipeline is compatible with any cell sorting experiment performed on a cellenONE instrument, ranging from cell line generation to other single-cell ‘omics (scRNAseq, etc). The pipeline has three main stages.

The first stage focuses on image cropping, segmentation and cell counting. For this task, the dispensed cell image is first cropped to a square surrounding the isolated cell, which is then processed using a trained Cellpose-SAM model to produce cell masks to finally determine singlet/multiplet dispensing.

In the second stage of cell phenotyping), the cropped images and cell masks are further processed by Cellprofiler. This permits further extraction of phenotypic information, over and above the diameter, elongation and circularity metrics offered routinely by the cellenONE. These additional features offer the opportunity to do either image-alone, or multi-omic classification and analysis paired with the SCP data. By extracting >200 additional metrics from cellenONE micrographs, our CellProfiler pipeline supplies a phenotypic layer that can be systematically aligned with proteomics data, enabling coherent multimodal characterisation of biological systems at single-cell resolution.

### A new module allows interactive exploration of cellenONE microscopy data associated with single-cell proteomics experiments using the ‘iSEE’ bioconductor package

The third and final part of our pipeline enables users to interactively explore and visualise both image-derived metadata and proteomics data prepared using *scpImaging* collectively in a customised panel within the ‘iSEE’ environment^21^. The design features a custom iSEE panel we term the ‘CellenONE plot’ (**Fig 7).** This panel shows both the unmodified image file and background file recorded during the run, as well as images showing the cell outlines and image masks generated using the *scpImaging* pipeline. It offers the ability for manual exploration and quality control of datasets and outliers. Single-cells can be selected for investigation in the app from other panels, for example dimensionality reduction plots, or tables of single-cells. For implementation, all cell images need to be placed within the ‘www’ subdirectory of the ‘iSEE’ app, to be accessible during its runtime. The HEK-293T/A549 dataset from this manuscript is available for online interactive data exploration at https://emmottlab.shinyapps.io/scpImaging_HEKA549/.

**Figure 7.**
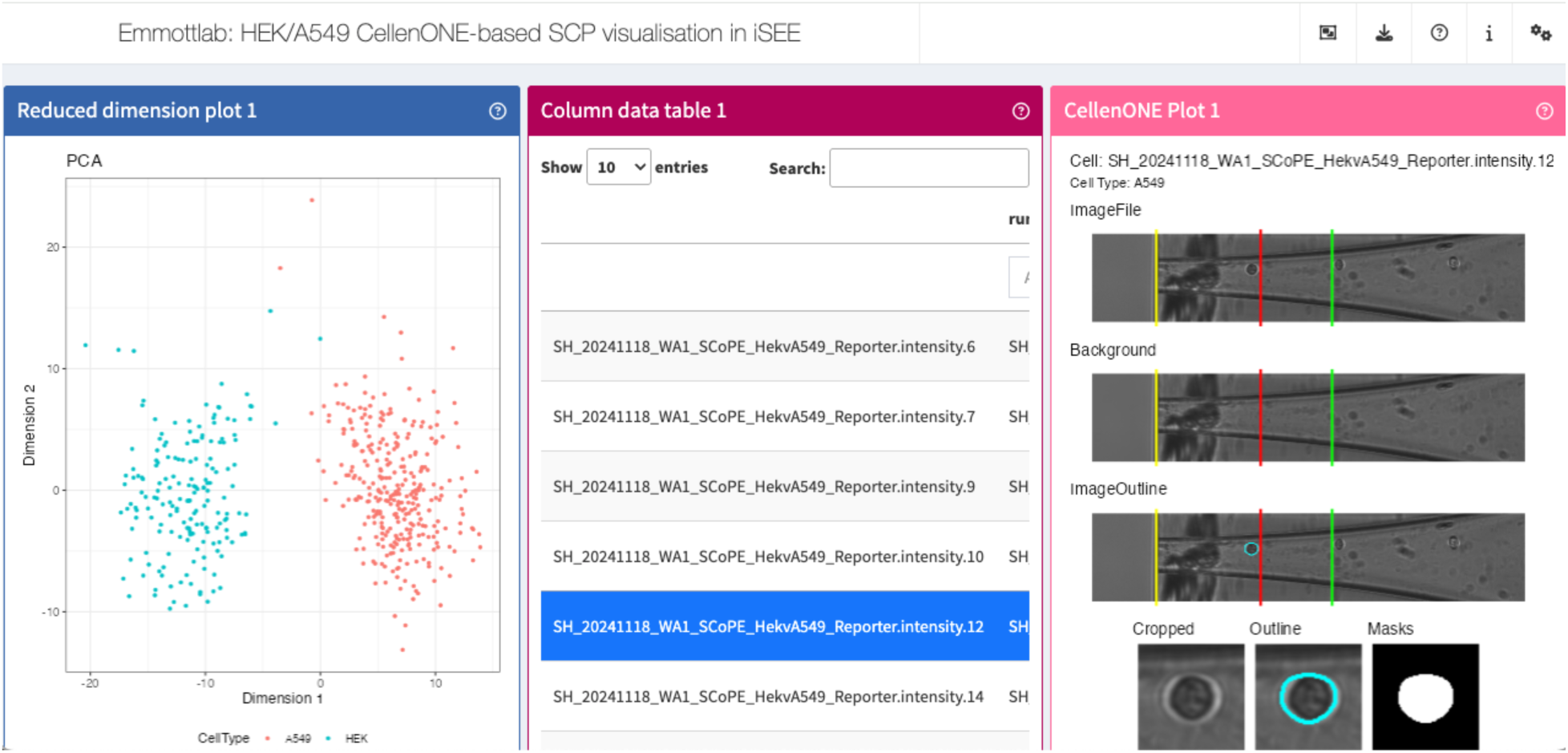
CellenONE-derived imaging data associated with single-cell experiments can be interactively explored through a custom iSEE panel. An ImageFile is recorded for each cell dispensed by the cellenONE. The yellow lines show the end of the PDC, red lines the border of the ejection zone, and the green lines the border of the sedimentation zone. Background images are produced by the cellenONE to determine background levels. The zoomed in image is a 75 x 75 pixel crop from the main image, centred on the cell coordinates provided by the cellenONE metadata, with the mask and outline version of the image generated in Cellpose and the ‘scpImaging’ R package. The CellenONEplot custom iSEE panel is included in the *scpImaging* package.

## Discussion

In the present study, we introduce a first approach for detection of doublets and multiplets in SCP samples prepared using the cellenONE system. As a principle, it relies on the advantageous nature of imaging-based analysis, which permits both automated and manual verification on a per-cell basis if required.

Doublets represent a well-recognised challenge for single-cell omics analysis and have been extensively investigated in single-cell transcriptomics. In scRNA-seq datasets, these artefacts can represent a range of events from 1-2% to nearly 40% of measured cells. Their occurrence is platform-dependent and operator choices in experimental design (e.g. cell concentrations) can further influx incidence^33^. Fluorescence-activated cell sorting (FACS) and cellenONE-based systems are commonly employed for single cell isolation, both of which utilise gating strategies to define single-cell events. Depending on the cells and biological question under analysis, more relaxed gating may be desired to capture a wider range of cell morphologies and thus biology. The stringency of these strategies introduces an inherent trade-off; stricter criteria may reduce multiplet capture but may exclude biologically relevant heterogeneity, whereas more permissive parameters retain that broader biological spectrum at the increased risk of multiplet inclusion. Even in cellenONE workflows which can achieve very low doublet rates, as reflected in our data, (**Fig. 1**) doublets can nevertheless arise and therefore warrant attention and detection.

Image-based verification as implemented in our *scpImaging* pipeline offers direct post-acquisition screening for doublets. A core advantage of our approach is that the images and data required are obtained during standard cellenONE-based preparations. Given that these micrographs and metadata are likely already available for majority of previously published cellenONE-based SCP datasets, *scpImaging* can be applied retrospectively without any modifications to the original experimental workflows. Our strategy is anchored in the cell dispensing step yet is agnostic to the type of sample preparation used downstream. Here, our data originate from nPOP sample preparations, but *scpImaging*’s applicability can extend to any cellenONE compatible method, such us EVO-96 based preparation, cellenCHIPs and even in clonal cell line generation or other cellenONE-based sc-omics. Although its general application is confined to the cellenONE platform, the conceptual foundation is broadly transferable, and the framework could be adapted in the future to other microscopy-based single-cell dispensing systems.

The ability to identify doublets naturally raises the question of their potential impact on downstream proteomics analysis. With respect to this experimental question, we constructed a controlled multiplexed TMT-based layout with defined empty, singlet and doublet containing droplets – greatly emphasising homotypic and heterotypic doublet cells. Of interest here, is that standard metrics (e.g. CV, number of peptides etc.) used to identify and remove poor-quality single cells fail at removing doublets, as they are designed to detect cells with low quantitative signal, poor quality quantitation, and very high sample to carrier ratios. Accordingly, doublets marginally score better compared to singlets reflecting their generally higher protein abundance. One area where our workflow performs less well is in detection of failed cell dispensing, as this capability sits outside its primary purpose; these cases are more effectively managed by existing QC filters.

These findings underscore the necessity for new and improved metrics or approaches to exclude these doublets in downstream analysis. Our pipeline embodies a solution to systematically identify and eliminate these artefacts, thereby preserving the integrity of SCP data. In line with observations from scRNAseq studies, heterotypic doublets here form an intermediate population, between the two parental populations, as this type of doublet capture averages protein-level expression between the two singlet populations. Homotypic doublets are a much more challenging population to remove computationally, by non-imaging-based doublet removal methods. In our data, we note homotypic cells showing slight separation within the singlet populations, adopting more extreme separation along PC1 than the singlets, remaining within the same cluster but towards the periphery. It is possible that the higher protein level results in differential ratio compression between the doublets and singlets. Our study which employed TMT-based analysis could therefore be inflating the level of separation observed for homotypic doublets, with ratio compression a feature of TMT-based labelling strategies. Analysis of label-free datasets in future work is encouraged to confirm if this separation is genuine or label-biased. However, the pattern observed with the periphery-skewed singlet cluster is consistent with prior observations from scRNA-seq data, so this observation could be authentic rather than artefactual^11, 14, 34^.

Despite the low-resolution of the captured images, we validated that cellenONE-based micrographs contain additional information that can be extracted using widely available imaging analysis software such as CellProfiler^20^. This imaging data alone permitted cell classification of our two-cell line dataset. More complex datasets involving a range of cell types or primary cells may be more challenging for classification but should be considered in follow up experiments. High classification AUCs were obtained when classifying cells based on the segmentation mask (cell area) portion of the image alone. Significant improvements in classification were recorded when using the entire image. It must be recognised that, since in our experimental design, the two cell lines were dispensed interchangeably, batch effect from sample build-up on the capillary, outside of our control, might have been introduced. For mixtures of different cell types, picked up and dispensed as a single population, use of the whole images may be a viable approach, though the time of dispensing, which is recorded by the cellenONE, should be considered as a covariate to control for changes in capillary appearance throughout the cell dispensing run.

In line with the cellenONE defaults, we have only recorded micrographs for acquired single cells. This default is used to reduce disk space requirements during sample acquisition, although the instrument can record all single cells observed during the run including those falling within or outside the acquisition gating settings. Future experimental designs could include significantly larger imaging components analysing all identified cell images, where a subset of the imaged cells also possess SCP data, rather than just cells prepared for single-cell omics, considering how sample throughput for SCP remains challenging. This would also permit post-acquisition analysis of important populations that have been excluded from the gated population analysed.

The latent utility of cellenONE-based microscopy data, and its corresponding metadata is fully exemplified by the phenotypic dimension it offers to single-cell analysis. At the same time, the results illuminate key limitations and areas for future improvement. The integrative analysis performed is limited in scope, correlating features to principal components. More extensive and complex analyses are certainly possible, though their reliability may be influenced by variation in image quality and cell focus. It is worth noting that the cellenONE microscope is designed to capture images of cells, with the focus on the capillary rather than the single cells, contributing to out of focus cell images. Technical advancements in control software could involve use of the fine x,y,z-axis controls of the cellenONE to automatically acquire in-focus images or z-stacks of identified cell images.

Summary information from fluorescent images in cases where this functionality is used are included in the data exported by the cellenONE, but we have not explored CellProfiler or Cellpose-SAM to further analyse these images. Our focus in this study has been primarily on the transmitted light images, but the cellenONE’s fluorescence sorting mode is a possible avenue for future experimentation, and additional iSEE panels or modifications to our cellenONE iSEE panel could be made to support visualisation of these images. Moreover, our pipeline has been successfully applied on both very clean and ‘dirty’ capillaries – the condition of these PDCs as well as any residual buildup can significantly impact image quality. Optimised selection of PDC coatings and chemistry may offer reduced build-up and improve image quality.

Our group has contributed to recent community guidelines on performing, benchmarking and reporting SCP data^35^. Thus, considering a range of datasets and instruments with *scpImaging* would be greatly beneficial for wider adaptation, standardisation and robustness of image-based doublet detection and analysis as part of SCP data. Finally, for reproducibility purposes and FAIR standards^36^, we would encourage cellenONE data sharing; including field files used, any generated micrographs, and log files during the cellenONE runs. As both SCP and spatial proteomics move towards more direct clinical applications^37^, the traceability offered by *scpImaging* for matching individual cell datapoints to images becomes even more crucial, both in terms of validation, and in the longer term for regulatory compliance.

To our knowledge, *scpImaging* represents the first SCP workflow capable of automated doublet detection using existing cellenONE metadata for QC, without adding experimental complexity or compromising throughput. Importantly, this does not require any modifications to the cellenONE instrument, or how samples are processed, as it relies on data already collected during standard operation. We have created an open-source R package and associated workflow, permitting other researchers to apply these methods to their own data and analysis. We further enhance analytical capacity by introducing a custom iSEE panel for interactive data exploration. The *scpImaging* pipeline is already positioned as a practical component for SCP and continued refinements in imaging fidelity, capillary chemistry and the investigation of diverse datasets will broaden its analytical scope.

## Acknowledgements

Edward Emmott is funded by an Academy of Medical Sciences Springboard Award (SBF006\1008) supported by the British Heart Foundation, Diabetes UK, the Global Challenges Research Fund, the Government Department for Business, Energy and Industrial Strategy and the Wellcome Trust. The Medical Research Council [MR/X000885/1], BBSRC [BB/W019744/1] and a Wellcome Trust Career Development Award [227831/Z/23/Z]. Trevor Sweeney is supported by a Wellcome Trust Sir Henry Dale Fellowship [202471/Z/16/A] and BBSRC grants to the Pirbright Institute [BBS/E/PI/230001A, BBS/E/PI/230002A, BBS/E/PI/23NB0003]. Jessica Coomber is funded by a joint Pirbright Institute-University of Liverpool PhD studentship. Markella Loi is supported by a BBSRC NW DLA CASE studentship in partnership with Cellenion. Stephen Holmes (2022) and Dr Leandro Neves (2023) have previously received Tandem Mass Tag Research Awards from Thermo Fisher Scientific in support of their research. Dr Maximilian Edrmann and Dr Frazer Buchanan (University of Liverpool), are thanked for their help with the Evos imager. Dr Rebekah Penrice-Randal for her comments on the draft manuscript. Cellenion, and Joshua Cantlon, as well as Andrew Leduc (Northeastern University) are thanked for their support with the nPOP protocol and cellenONE instrument.

## Author contributions

**ML and SH** prepared samples, performed Cellpose analysis, developed and tested pipelines, and provided supervision**, SZ** and **JC** performed Cellpose analysis. **MH** developed initial CellProfiler pipelines**, LXN** and **PJB** supported mass spectrometry sample preparation, acquisition and analysis**. EE** generated preliminary data, and performed data analysis and package development. **EE** and **TS** provided funding and supervision. All authors helped prepare the manuscript. The final version was read and approved by all authors.

## Supplementary material

Supplementary material includes supplementary figures S1-S7.

## Supplementary Figures

**Figure S1.**
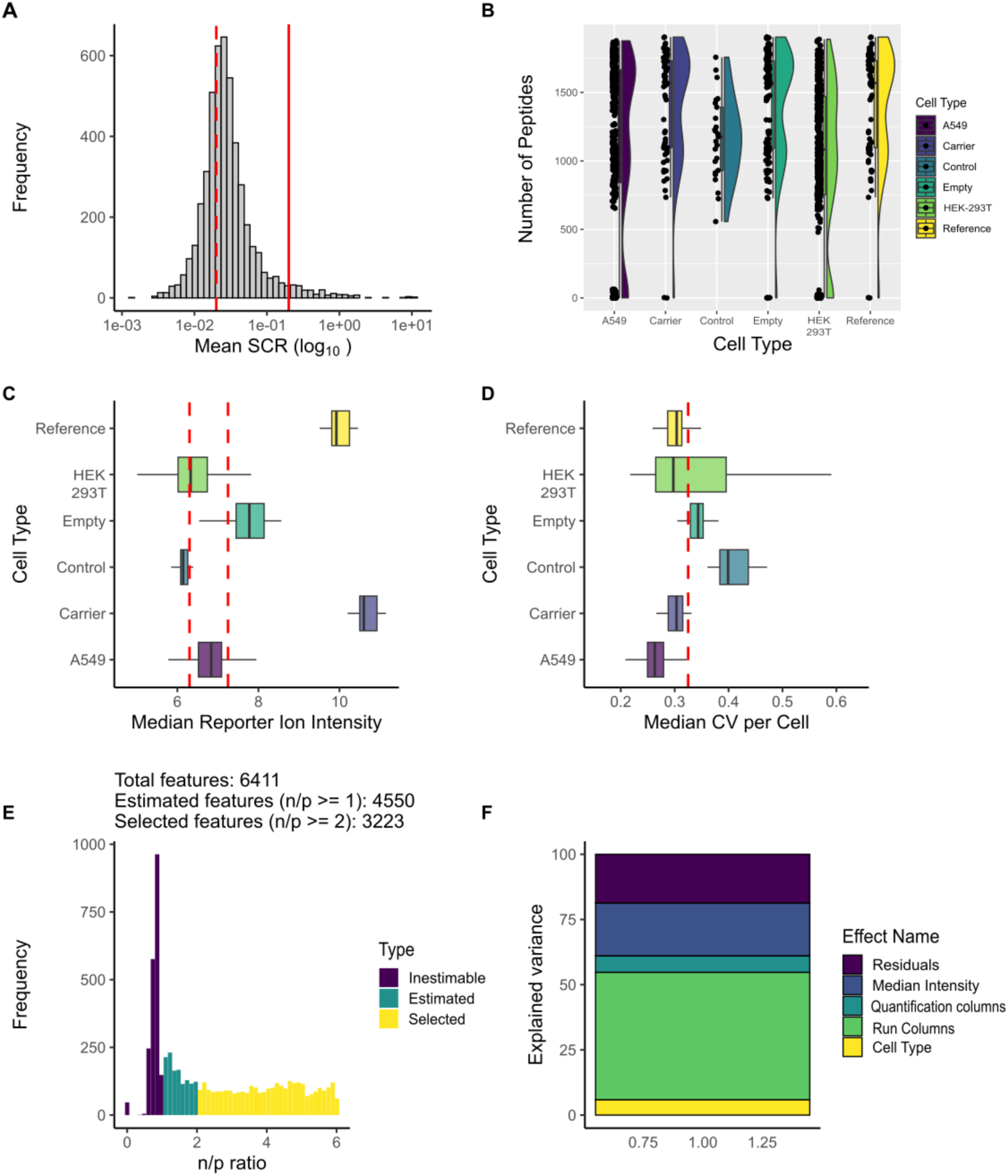
Single-cell proteomic data processing and modelling QC for the initial HEK-293T/A549 dataset. Data were processing using the scp Bioconductor package, using a minimal data processing workflow.Samples were filtered based on a) sample-to-carrier ratio, b) number of peptides, c) Median reporter ion intensity, and d) median CV per cell. Cutoffs applied to the data are indicated with red lines. For linear modeling using scplainer, a e) n/p ratio of >= 2 was used to select features. The variance explained for each effect in the model is given in f).

**Figure S2.**
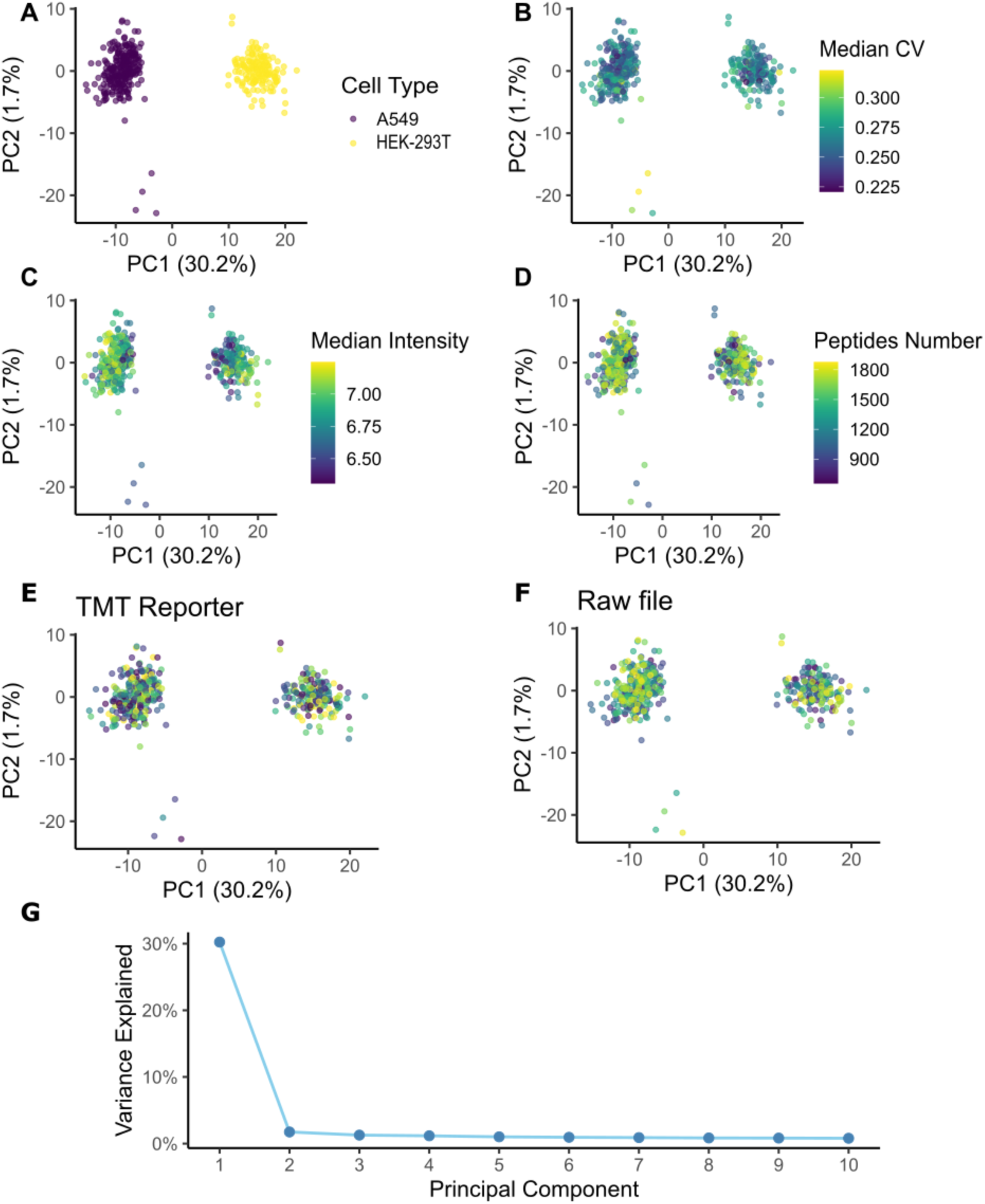
Single-cell proteomics data QC for the initial HEK-293T/A549 dataset. A) HEK-293T and A549 cells separate cleanly by cell type along PC1. A few cells cluster beneath the main A549 cluster and have higher B) CV, and C) lower median intensity, and lower numbers of D) identified peptides, suggest sample preparation issues for those four cells. Overlaying TMT reporter or Raw file (batch) shows the absence of batch effects. PC1 and 2 explain the most variance in this dataset with PC1 dominating.

**Figure S3.**
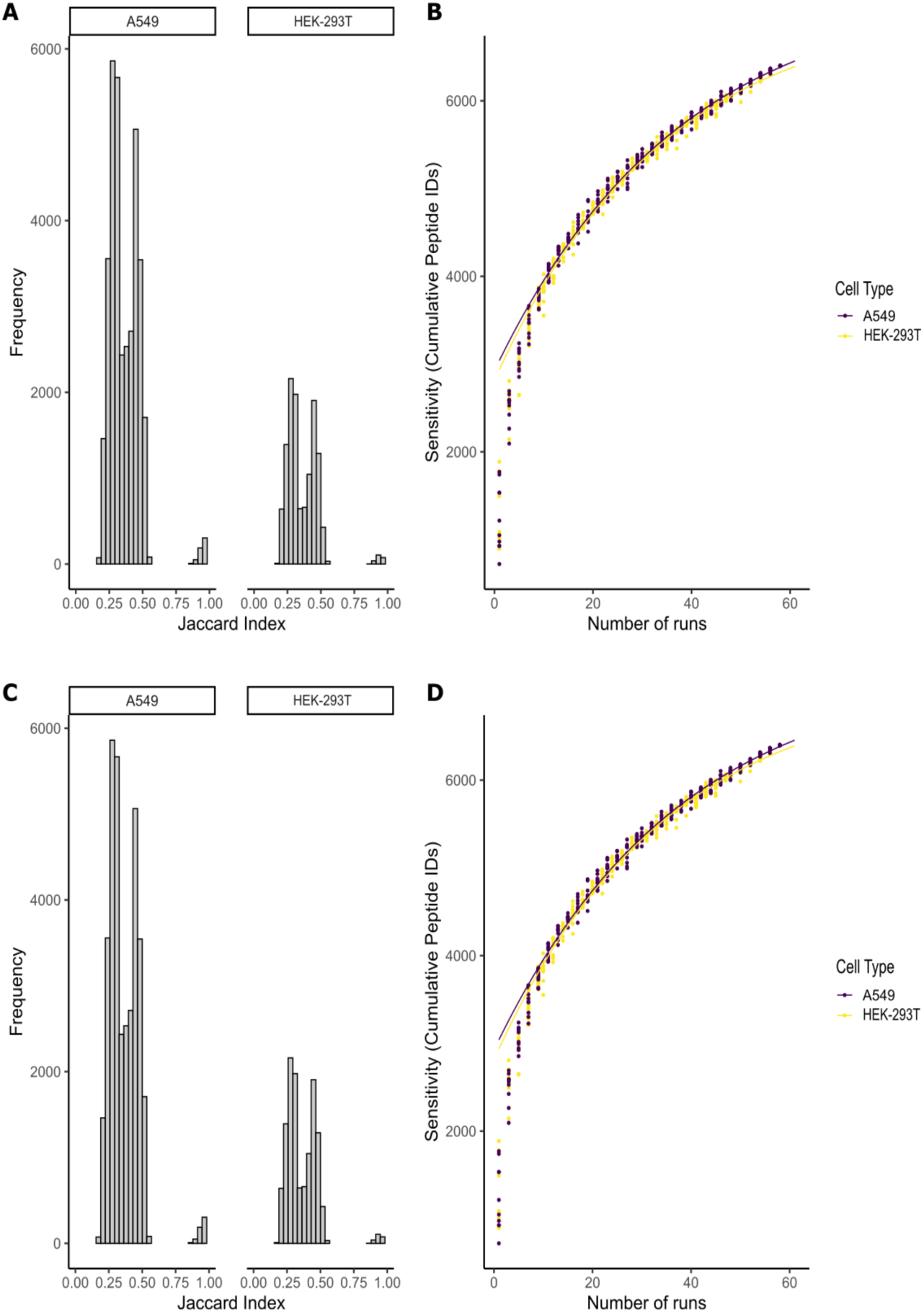
Single-cell proteomics reporting data for the initial HEK-293T/A549 cell dataset. A) Overlap (Jaccard index) in data measured between HEK-293T and A549 cells. B) Cumulative sensitivity curve illustrating total peptide coverage for this dataset. C) Overlap in protein data measured between HEK-293T and A549 cells. D) Cumulative sensitivity curve illustrating total protein coverage for this dataset.

**Figure S4.**
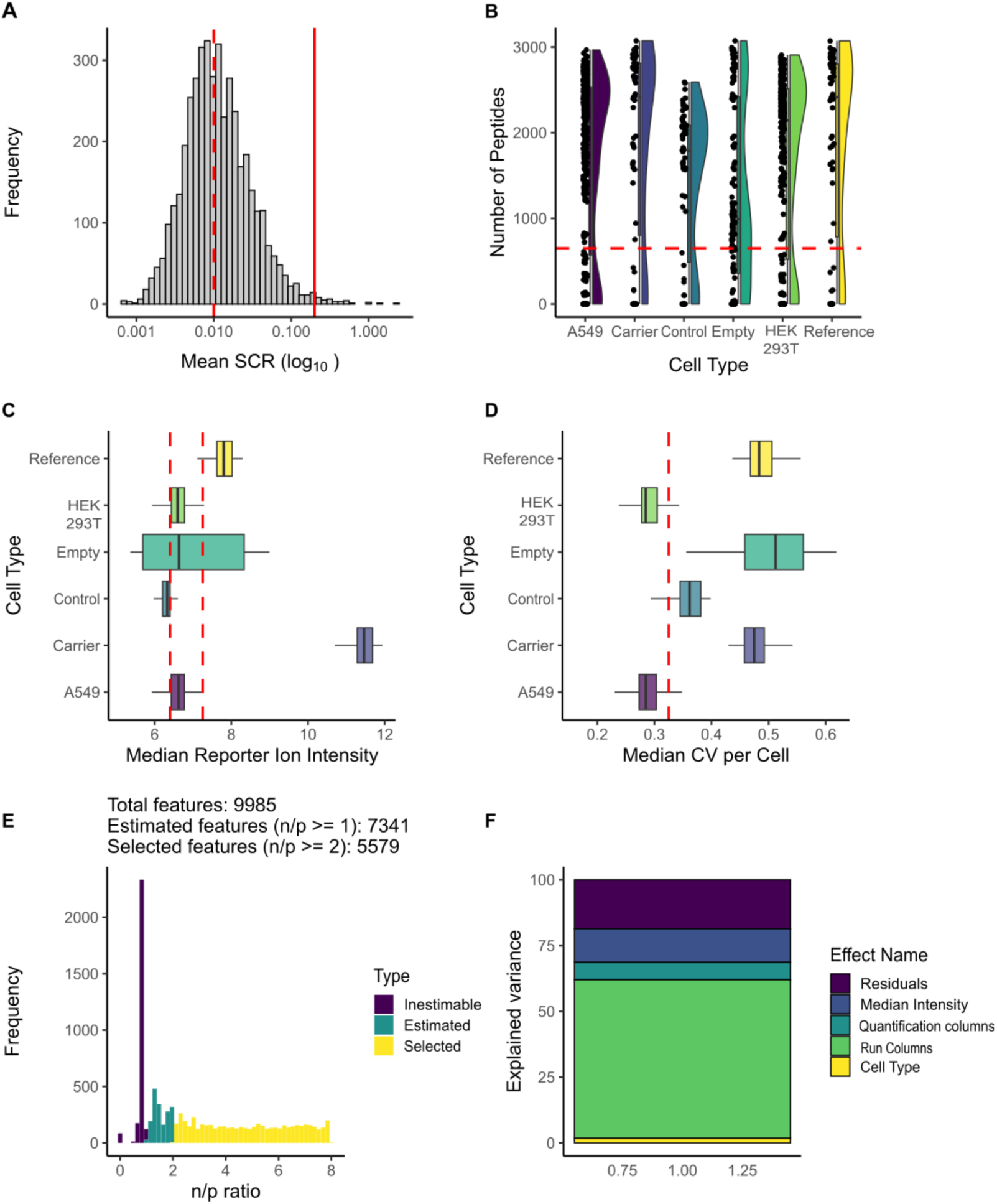
Single-cell proteomic data processing and modelling QC for the HEK-293T/A549 singlet/doublet dataset. Data were processing using the **scp** Bioconductor package, using a minimal data processing workflow. Samples were filtered based on a) sample-to-carrier ratio, b) number of peptides, c) Median reporter ion intensity, and d) median CV per cell. Cutoffs applied to the data are indicated with red lines. For linear modeling using scplainer, a e) n/p ratio of >= 2 was used to select features. The variance explained for each effect in the model is given in f).

**Figure S5.**
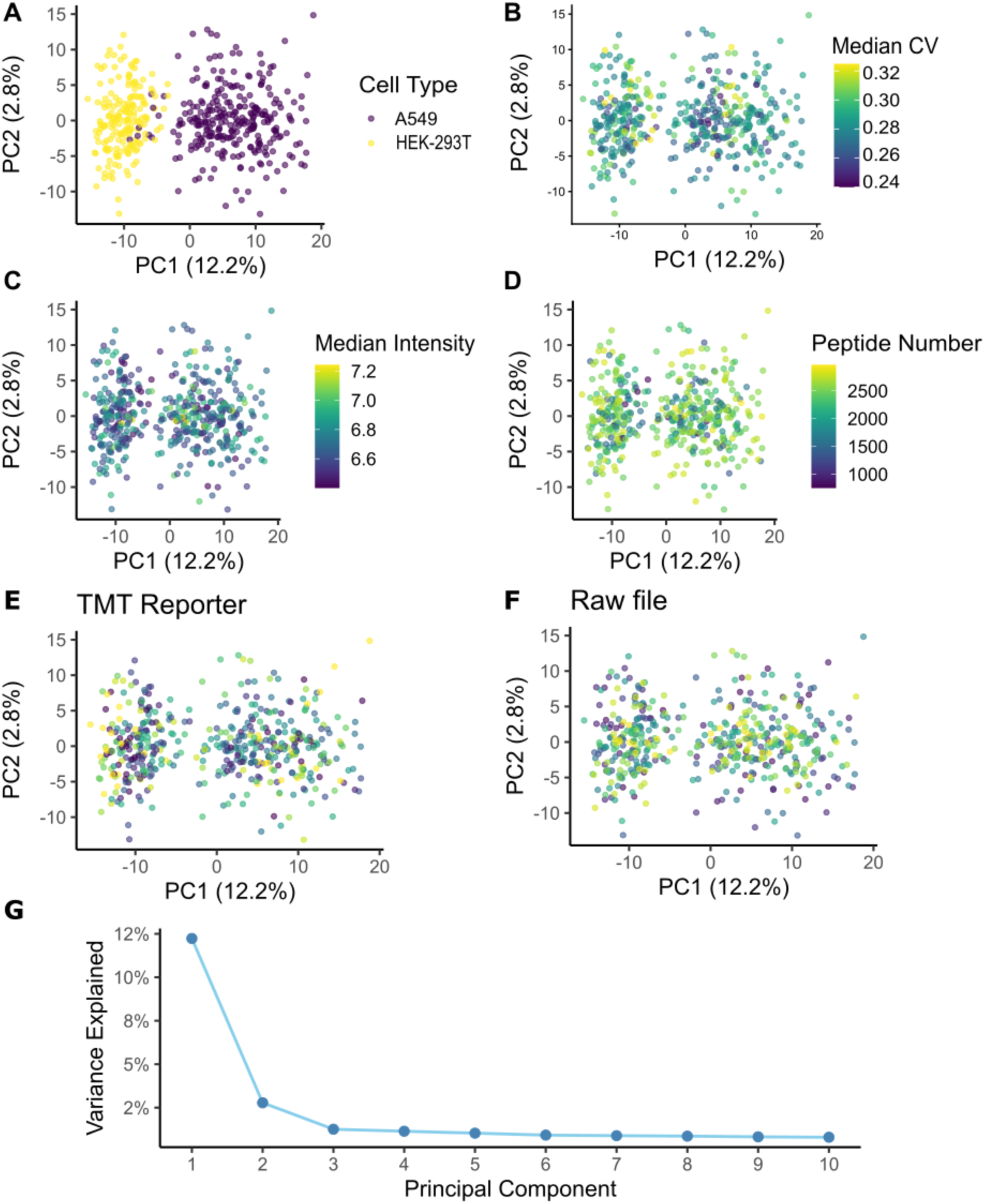
Single-cell proteomics data QC for the HEK-293T/A549 singlet/doublet dataset. HEK-293T and A549 cells separate cleanly by cell type along PC1. B) shows the median CV, C) median peptide intensity, D) Number of identified peptides. Overlaying E) TMT reporter or F) Raw file (batch) shows the absence of batch effects. G) PC1 and 2 explain the most variance in the dataset, with PC1 dominating.

**Figure S6.**
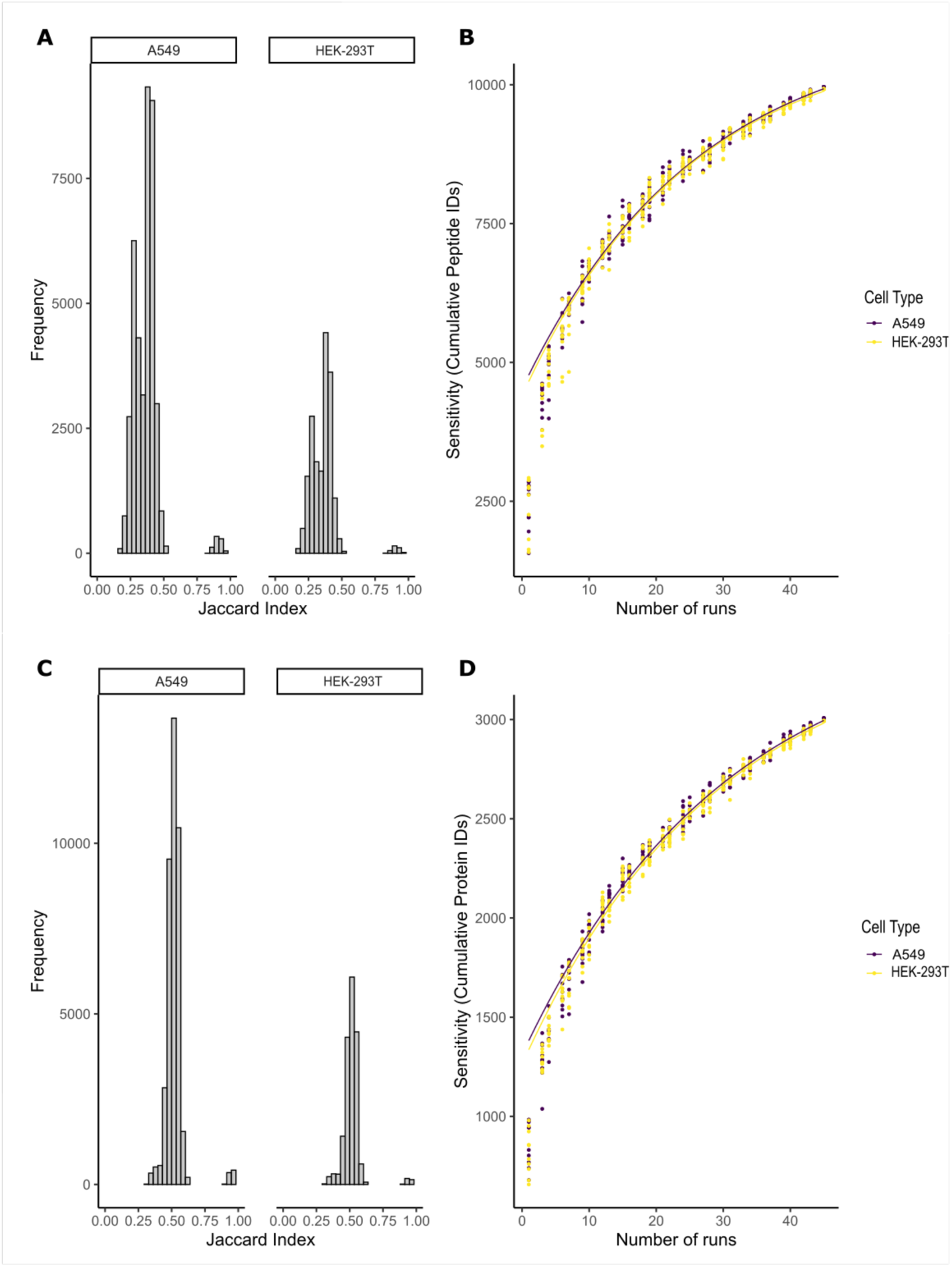
Single-cell proteomics reporting data for the HEK-293T/A549 singlet/doublet experiment. A) Overlap (Jaccard index) in peptide data measured between HEK-293T and A549 cells. B) Cumulative sensitivity curve illustrating total peptide coverage for this dataset. C) Overlap in protein data measured between HEK-293T and A549 cells. D) Cumulative sensitivity curve illustrating total protein coverage for this dataset.

**Figure S7.**
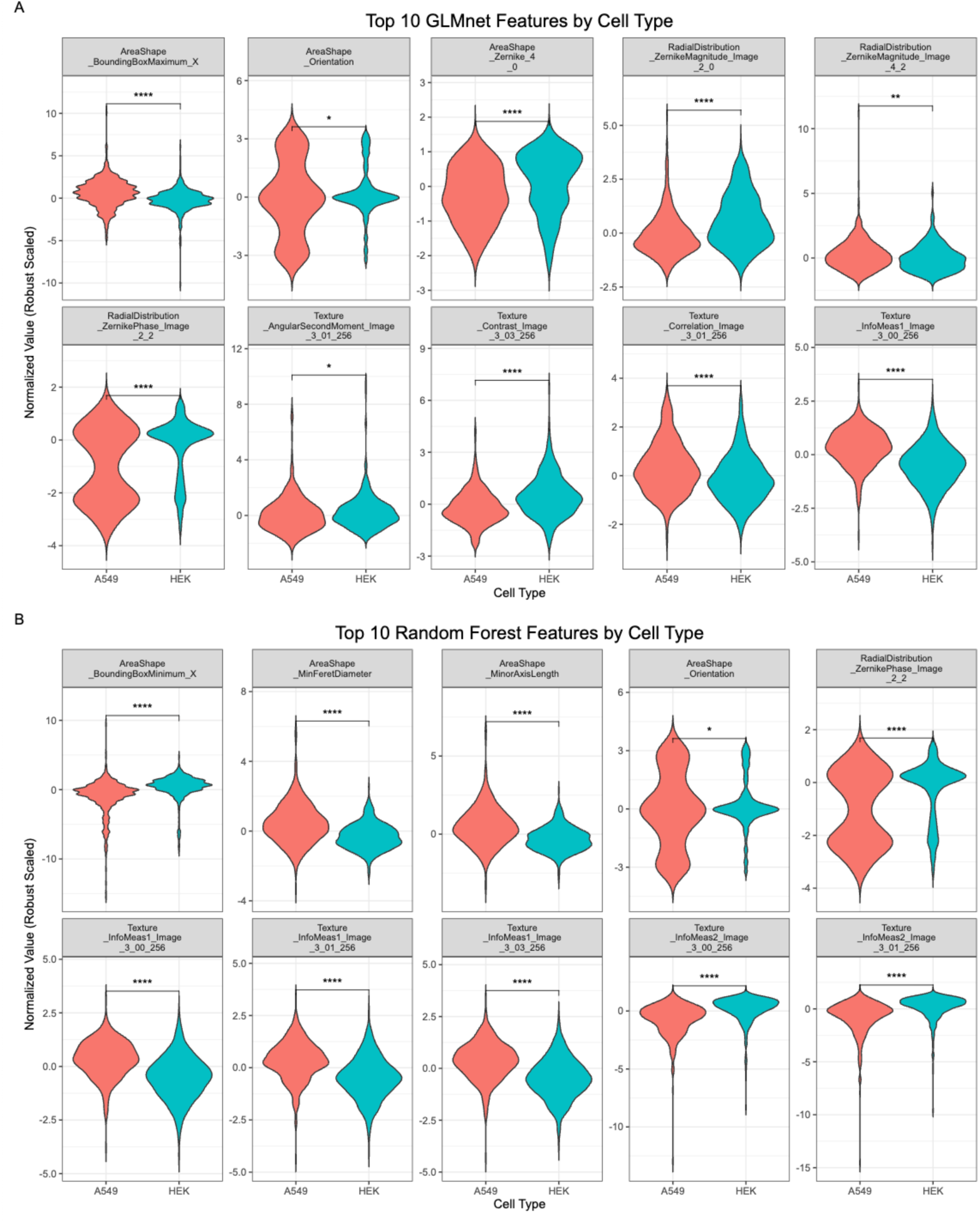
Top 10 Cellprofiler classifier features for GLM and Random Forest-based classifiers based on segmentation masks. Note, the Top ten features do show overlap between these models (e.g. AreaShape_Orientation), and we have chosen to show all ten features for both models which results in a subset of panels displaying for both a) the GLM-based model and b) the Random Forest model.

## References

1. Hartlmayr D, Ctortecka C, Mayer R, Mechtler K, Seth A. Label-Free Sample Preparation for Single-Cell Proteomics. In: Mass Spectrometry Based Single Cell Proteomics (eds Vegvari A, Teppo J, Zubarev RA). Springer US (2024).

2. Altschuler SJ, Wu LF. Cellular Heterogeneity: Do Differences Make a Difference? Cell 141, 559–563 (2010).

3. Gupta RK, Kuznicki J. Biological and Medical Importance of Cellular Heterogeneity Deciphered by Single-Cell RNA Sequencing. Cells 9, (2020).

4. Adams Sophia K, Ducharme Grace E, Loveday Emma K. All the single cells: if you like it then you should put some virus on it. Journal of Virology 98, e01273–01223 (2024).

5. Phillip JM, Han K-S, Chen W-C, Wirtz D, Wu P-H. A robust unsupervised machine-learning method to quantify the morphological heterogeneity of cells and nuclei. Nature Protocols 16, 754–774 (2021).

6. Goldman SL, MacKay M, Afshinnekoo E, Melnick AM, Wu S, Mason CE. The Impact of Heterogeneity on Single-Cell Sequencing. Frontiers in Genetics Volume 10 - 2019, (2019).

7. Hasin Y, Seldin M, Lusis A. Multi-omics approaches to disease. Genome Biology 18, 83 (2017).

8. Budnik B, Levy E, Harmange G, Slavov N. SCoPE-MS: mass spectrometry of single mammalian cells quantifies proteome heterogeneity during cell differentiation. Genome Biology 19, 161 (2018).

9. Petelski AA, et al. Multiplexed single-cell proteomics using SCoPE2. Nat Protoc 16, 5398–5425 (2021).

10. Kang HM, et al. Multiplexed droplet single-cell RNA-sequencing using natural genetic variation. Nature Biotechnology 36, 89–94 (2018).

11. Wolock SL, Lopez R, Klein AM. Scrublet: Computational Identification of Cell Doublets in Single-Cell Transcriptomic Data. Cell Systems 8, 281–291.e289 (2019).

12. Bais AS, Kostka D. scds: computational annotation of doublets in single-cell RNA sequencing data. Bioinformatics 36, 1150–1158 (2020).

13. DePasquale EAK, et al. DoubletDecon: Deconvoluting Doublets from Single-Cell RNA-Sequencing Data. Cell Reports 29, 1718–1727.e1718 (2019).

14. Luecken MD, Theis FJ. Current best practices in single-cell RNA-seq analysis: a tutorial. Molecular Systems Biology 15, e8746 (2019).

15. Xi NM, Li JJ. Benchmarking Computational Doublet-Detection Methods for Single-Cell RNA Sequencing Data. Cell Systems 12, 176–194.e176 (2021).

16. Vanderaa C, Gatto L. Revisiting the Thorny Issue of Missing Values in Single-Cell Proteomics. Journal of Proteome Research 22, 2775–2784 (2023).

17. Lin H-JL, Webber KGI, Nwosu AJ, Kelly RT. Review and Practical Guide for Getting Started With Single-Cell Proteomics. PROTEOMICS 25, e202400021 (2025).

18. Matzinger M, Mayer RL, Mechtler K. Label-free single cell proteomics utilizing ultrafast LC and MS instrumentation: A valuable complementary technique to multiplexing. PROTEOMICS 23, 2200162 (2023).

19. Stringer C, Wang T, Michaelos M, Pachitariu M. Cellpose: a generalist algorithm for cellular segmentation. Nature Methods 18, 100–106 (2021).

20. Stirling DR, Swain-Bowden MJ, Lucas AM, Carpenter AE, Cimini BA, Goodman A. CellProfiler 4: improvements in speed, utility and usability. BMC Bioinformatics 22, 433 (2021).

21. Rue-Albrecht K, Marini F, Soneson C, Lun ATL. iSEE: Interactive SummarizedExperiment Explorer. F1000Res 7, 741 (2018).

22. Leduc A, Khoury L, Cantlon J, Khan S, Slavov N. Massively parallel sample preparation for multiplexed single-cell proteomics using nPOP. Nature Protocols 19, 3750–3776 (2024).

23. Nie S, O’Brien Johnson R, Livson Y, Greer T, Zheng X, Li N. Maximizing hydrophobic peptide recovery in proteomics and antibody development using a mass spectrometry compatible surfactant. Analytical Biochemistry 658, 114924 (2022).

24. Tyanova S, Temu T, Cox J. The MaxQuant computational platform for mass spectrometry-based shotgun proteomics. Nature Protocols 11, 2301–2319 (2016).

25. Chen AT, Franks A, Slavov N. DART-ID increases single-cell proteome coverage. PLoS Comput Biol 15, e1007082 (2019).

26. Pachitariu M, Rariden M, Stringer C. Cellpose-SAM: superhuman generalization for cellular segmentation. bioRxiv, 2025.2004.2028.651001 (2025).

27. Carpenter AE, et al. CellProfiler: image analysis software for identifying and quantifying cell phenotypes. Genome Biology 7, R100 (2006).

28. Vanderaa C, Gatto L. scplainer: using linear models to understand mass spectrometry-based single-cell proteomics data. Genome Biology 26, 237 (2025).

29. Vanderaa C, Gatto L. Replication of single-cell proteomics data reveals important computational challenges. Expert Rev Proteomics 18, 835–843 (2021).

30. Vanderaa C, Gatto L. The Current State of Single-Cell Proteomics Data Analysis. Curr Protoc 3, e658 (2023).

31. Gurazada SGR, Kennedy HM, Braatz RD, Mehrman SJ, Polson SW, Rombel IT. HEK-omics: The promise of omics to optimize HEK293 for recombinant adeno-associated virus (rAAV) gene therapy manufacturing. Biotechnology Advances 79, 108506 (2025).

32. Ujie M, et al. Long-term culture of human lung adenocarcinoma A549 cells enhances the replication of human influenza A viruses. Journal of General Virology 100, 1345–1349 (2019).

33. Ttoouli D, Hoffmann D. Multiplets in scRNA-seq data: Extent of the problem and efficacy of methods for removal. PLOS ONE 20, e0333687 (2025).

34. McGinnis CS, Murrow LM, Gartner ZJ. DoubletFinder: Doublet Detection in Single-Cell RNA Sequencing Data Using Artificial Nearest Neighbors. Cell Systems 8, 329–337.e324 (2019).

35. Gatto L, et al. Initial recommendations for performing, benchmarking and reporting single-cell proteomics experiments. Nat Methods 20, 375–386 (2023).

36. Go FAIR.) (2025).

37. Nordmann TM, et al. Spatial proteomics identifies JAKi as treatment for a lethal skin disease. Nature 635, 1001–1009 (2024).

